# Evolution during seed production for ecological restoration? A molecular analysis of 19 species finds only minor genomic changes

**DOI:** 10.1101/2021.11.03.467064

**Authors:** Malte Conrady, Christian Lampei, Oliver Bossdorf, Walter Durka, Anna Bucharova

## Abstract

A growing number of restoration projects require large amounts of seeds. As harvesting natural populations cannot cover the demand, wild plants are often propagated in large-scale monocultures. There are concerns that this cultivation process may cause genetic drift and unintended selection, which would alter the genetic properties of the cultivated populations and reduce their genetic diversity. Such changes could reduce the pre-existing adaptation of restored populations, and limit their adaptability to environmental change.

We used single nucleotide polymorphism (SNP) markers and a pool-sequencing approach to test for genetic differentiation and changes in gene diversity during cultivation in 19 wild grassland species, comparing the source populations and up to four consecutive cultivation generations grown from these sources. We then linked the magnitudes of genetic changes to the species’ breeding systems and seed dormancy, to understand the roles of these traits in genetic change.

The propagation of native seeds for ecosystem restoration changed the genetic composition of the cultivated generations only moderately. The genetic differentiation we observed as a consequence of cultivation was much lower than the natural genetic differentiation between different source regions, and the propagated generations harbored even higher gene diversity than wild-collected seeds. Genetic change was stronger in self-compatible species, probably as a result of increased outcrossing in the monocultures.

**Synthesis and applications:** Our study indicates that large-scale seed production maintains the genetic integrity of natural populations. Increased genetic diversity may even increase the adaptive potential of propagated seeds, which makes them especially suitable for ecological restoration. However, we have been working with seeds from Germany and Austria, where the seed production is regulated and certified. Whether other seed production systems perform equally well remains to be tested.

## Introduction

Ecological restoration of degraded habitats is an indispensable tool for handling the current biodiversity crisis (www.decadeonrestoration.org; Díaz et al. 2019). Degraded terrestrial ecosystems frequently lack diaspores from which new communities could regenerate and successful restoration therefore often requires introduction of seeds from other sources (Hölzel et al. 2012; Elzenga et al. 2019). With upscaling restoration, the demand for native seeds is increasing and the shortage of seed has become one of the major obstacles of restoration projects (Merritt & Dixon 2011; Nevill et al. 2018). Harvesting seeds from the wild is not sustainable because seeds are the basic means of plant reproduction and their excessive removal may threaten the persistence of source populations (Meissen et al. 2015). Consequently, wild-collected seeds are often propagated on farms to increase their amounts, and farm-produced seeds are then used for restoration projects (Kiehl et al. 2014; Nevill et al. 2018; Bucharova et al. 2019)

Agricultural propagation of seeds sourced from wild population aims to maintain the natural integrity of the collected seed (Espeland et al. 2017; Bucharova et al. 2019). This differs from native plant cultivars which are commonly bred to obtain specific characteristics like rapid growth, seedling vigour or high seed production (Leger & Baughman 2015). Yet, even seed propagation that does not intentionally select certain genotypes could affect the genetic integrity of cultivated populations because the propagation process may cause genetic drift or unintended selection (Espeland et al. 2017). In the seed beds, plants are often grown in monocultures, with additional watering, fertilization and sometimes protection from herbivores, releasing the plants from natural selection by these factors. Machine harvesting and seed cleaning can further select for seeds of specific characteristics like shape or weight (Espeland et al. 2017). The seeds are often propagated for multiple generations, with a subset of cultivated seeds used for the establishment of the next cultivated generation (Bucharova et al. 2019). This may result in repeated genetic bottlenecks and genetic drift, i.e. random changes in allele frequency and loss of genetic variability. Populations may also randomly accumulate alleles that reduce adaptation to natural environments (Pertoldi et al. 2007; Lau et al. 2019). The loss of genetic variability would be particularly problematic because standing genetic variation is a necessary prerequisite for rapid adaptation (Crowe & Parker 2008). Restored populations with low genetic variability would have a reduced ability to adapt to changing conditions, for example under climate change (Barrett & Schluter 2008).

The severity of propagation effects on the genetic properties of a population will depend on the cultivation methods and life-history traits of the plants. For example, manual seed harvest throughout the season likely causes smaller changes than mechanical harvest because it samples a greater variety of phenologies and thus a larger proportion of genetic variability. The cultivation effects are further expected to decrease with increasing size of the cultivated population, because genetic drift is stronger in small populations (Frankham et al. 2014). Regarding species life-history traits, species with strong seed dormancy will likely change more during cultivation, because genotypes with dormant seeds will contribute less to the gene pool of the next generation (Kettenring & Galatowitsch 2007). Moreover, self-compatible species will likely change more because selection can act faster on predominantly self-pollinating species (Andersson & Ofori 2013). Indeed, one of the few existing studies on cultivation effects in plants propagated for restoration (Nagel et al. 2019) found the strongest genetic changes in a self-pollinating species.

Although genetic changes during cultivation are expected and intensely debated in the restoration literature (e.g. Basey et al. 2015; Espeland et al. 2017; Pedrini et al. 2020), there is surprisingly little empirical evidence for them. A substantial body of literature describes crop domestication (reviewed in Pickersgill 2009; Kantar et al. 2017), but in contrast to crops, plant materials cultivated for restoration are not intentionally bred. Other studies documented evolutionary changes in ex situ cultivation in botanical gardens (Ensslin & Godefroid 2019; Rauschkolb et al. 2019), but ex-situ populations are usually very small, whereas the cultivation of plant material for ecosystem restoration usually involves large populations spanning over many square meters to hectares. The few studies that focused specifically on evolution of plants for ecological restoration worked with individual species and/or documented cultivation only across a single generation (Dyer et al. 2016; Basey St. Clair et al. 2020; Pizza et al. 2021). Only one previous study focused on multiple cultivation generations across multiple species, and it found that cultivation effects differ between species and often appear to be minor (Nagel et al. 2019).

In this study, we used molecular markers to study the effect of agricultural propagation on the genetic variation of 19 different species of wild plants. We used seeds propagated in the German (Bucharova et al. 2019) and Austrian (Krautzer et al. 2020) system of seed production, where seeds collected from one or several wild populations are mixed and then propagated for up to five consecutive generations. For each species, we obtained wild-collected seeds and seeds from the one to four consecutive cultivation generations (Figure 1). We used genotyping-by-sequencing to assess the effect of cultivation on the genetic properties of the populations. We hypothesized that 1) the genetic divergence from the wild population will increase with the number of generations spent in cultivation, 2) the genetic diversity is decreasing with the number of generations spent in cultivation, and 3) that species characteristics predict part of the genetic changes experienced during cultivation.

**Figure 1:**
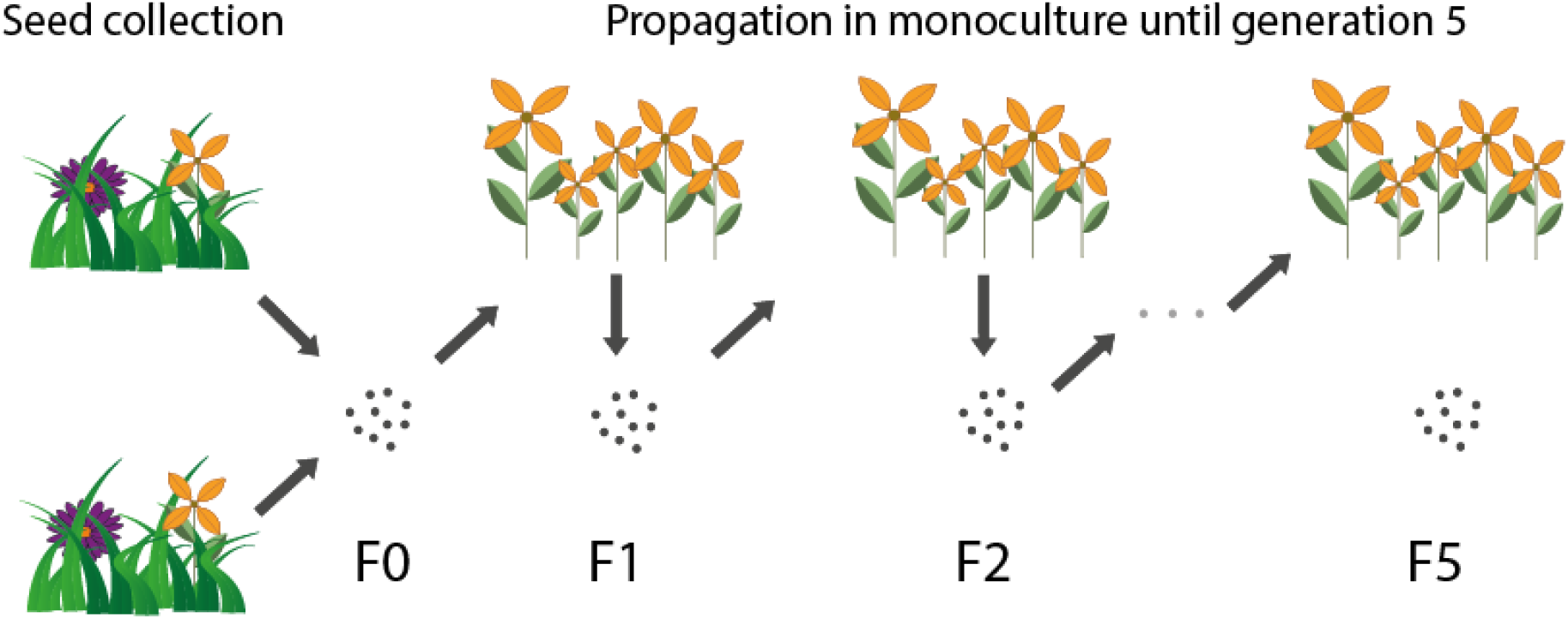
Schematic of the propagation process of wild plant seed material for restoration. Wild seeds are collected from multiple populations and mixed (F0). The F0 seeds are used to establish the first generation in cultivation (F1). These cultivated seeds are available for restoration projects, and a small part is used to establish the next cultivated generation. After five cultivated generations (F5), new seeds must be collected from the wild.

## Methods

### Seed material

We obtained seeds of grassland plants from two producers of seeds for ecological restoration, one in Austria and one in Germany, which we call Producer 1 and Producer 2 in the remainder of this paper. The seed propagation process starts with the collection of seeds in the wild. The Producer 1 bases its propagation on a mixture of seeds from multiple wild populations, whereas the Producer 2 on seeds coming from only one population. The wild-collected seeds (F0) are often first germinated in a greenhouse to produce plugs which are then planted into an agriculture field. The seeds of this first cultivated generation (F1) are then harvested and used to establish the next cultivation (F2, Figure 1). The F2 seeds are mostly sold, but some are kept to establish the F3 generation. This process is repeated until F5, then new seeds must be collected from the wild, to avoid that plants (presumably) adapt to their propagation environments or lose their genetic diversity (Figure 1) (Espeland et al. 2017). The seeds are usually mechanically harvested, using agricultural machinery.

For our study, we obtained wild-collected and cultivated seeds of 19 different plant species. Because the seed producers carefully stored the seeds from both the wild collection and almost every consecutive generation in cultivation, we were able to test for possible genetic changes during this cultivation process from generation to generation up to the fourth cultivated generation. Four of the species were provided by both producers, and one from two regions from the same producer (Table 1). In total we obtained 24 independent cultivation lines (19 species, 5 of them from two regions) which resulted in 83 accessions. To obtain material for the genetic analysis, we sowed seeds on seeding substrate, and sampled leaves from 18 random individuals per species and generation when the plants had become large enough.

**Table 1:** Species and generations in cultivation included in the experiment. F0 are wild collected seeds and F1-F4 are consecutive generations in cultivation. The generations in brackets were excluded from the data analyses because of insufficient SNP data and/or sampling depth (see text).

| Species | Family | Producer 1 | Producer 2 |
| --- | --- | --- | --- |
| *Achillea millefolium* | Asteraceae | F0-F2 | F0; F3; F4 |
| *Anthoxanthum odoratum* | Poaceae |  | F0-F2 |
| *Centaurea jacea* | Asteraceae | F0-F3 |  |
| *Centaurea scabiosa* | Asteraceae | F0-F2 | F0; F1; (F2); F3 |
| *Crepis biennis* | Asteraceae | F0-F2; (F3) |  |
| *Cynosurus cristatus* | Poaceae |  | F0-F2 |
| *Dianthus carthusianorum* | Caryophyllaceae |  | F0-F4 |
| *Galium verum* | Rubiaceae |  | F0-F2 |
| *Leontodon hispidus* | Asteraceae | F0-F2 |  |
| *Leucanthemum ircutianum* | Asteraceae | F0-F3 | F0-F2; F4 |
| *Lotus corniculatus* | Fabaceae | F0-F3 | F0; F1 |
| *Lychnis flos-cuculi* | Caryophyllaceae |  | F0-F2 |
| *Ranunculus acris* | Ranunculaceae | F0-F3 |  |
| *Rumex acetosa region R6* | Polygonaceae | F0; (F1); F2 |  |
| *Rumex acetosa region R16* | Polygonaceae | F0-F2 |  |
| *Salvia pratensis* | Lamiaceae |  | F0; F1; F3 |
| *Silene dioica* | Caryophyllaceae | F0-F3 |  |
| *Silene vulgaris* | Caryophyllaceae | F0-F3 |  |
| *Trifolium pratense* | Fabaceae | F0-F3 |  |
| *Veronica teucrium* | Plantaginaceae |  | F0-F2 |

### Molecular Analysis

Because of the large sample size, we used a population pool approach (Futschik & Schlötterer 2010), where we pooled 18 individuals per generation and cultivation line into one sample. We used a reduced-representation sequencing approach for SNP (single nucleotide polymorphism) detection and genotyping and followed the ddRAD protocol (Peterson et al. 2012) with slight deviations (see Supplementary Information).

We used process_radtags from the Stacks 2.0 pipeline (Catchen et al. 2013; Rochette et al. 2019) to demultiplex reads. We then used the dDocent 2.6.0 pipeline (Puritz et al. 2014; O’Leary et al. 2018) for contig assembly, SNP detection and assessment of allelic read counts for each cultivation line. SNP filtering of the resulting VCF file with vcftools removed indels, kept only biallelic loci with minimum Phred-scores of 30 and kept only one SNP per contig. To allow a comparison of genetic diversity between generations that is not biased by different sequencing depths, we further filtered the data using proprietary R scripts. We used only markers with a minor allele frequency of at least 0.05, and genotypes that had a minimum read depth of 36. We corrected for unequal sequencing depth of the same contig at different pools of the same cultivation line by rarefaction. After removing three pools with less than 500 SNPs (Tab. 1), the final data sets consisted of 657 - 9721 (average 5137) biallelic SNP loci per pool across the 19 species and 24 cultivation lines, with 0% to 14.6% missing data (Tab. S2). For analyses that compared different cultivation lines of the same species, the rarefaction procedure was repeated per species and not per cultivation line (See Supplementary information for details).

### Data analysis

All analyses were performed in R (R Development Core Team, 2020). First, we tested whether the genetic differentiation between the wild-collected and cultivated plants increased with the number of generations the population spent in cultivation. We calculated the pairwise genetic differentiation *F*_ST_ for each cultivation line between the plants from the wild-collected seeds and each consecutive generation in cultivation using the R package *poolfstat* (Hivert et al. 2018). We then centred the *F*_ST_ within each cultivation line by subtracting from each *F*_ST_ value the mean *F*_ST_ of that cultivation line, and related it to the generation number (continuous variable), producer identity and their interaction as explanatory variables and cultivation line as random factor in a linear mixed effects model using the package *lme4* (Bates et al. 2015). Only cultivation lines with three or more generations were included, because a minimum of two *F*_ST_ values was needed to centre around the mean (two cultivation lines excluded, see Table1).

To provide context for the magnitude of genetic differentiation caused by the cultivation process, we compared magnitude of genetic differentiation through cultivation to natural genetic differentiation between populations. Specifically, we compared the absolute values of genetic differentiation (*F*_ST_) between the wild plants and the last cultivated generation within a cultivation line with the *F*_ST_ between wild plants of the same species coming from two different regions using a linear mixed model (Bates et al. 2015) with *F*_ST_ as a response variable, wild vs. cultivated population as an explanatory variable and species as a random factor. This was possible in five species for which the F0 was available from two different regions, either from two seed producers or, in one species, from two regions from the same producer (Table 1). The geographical distance between the regions ranged from 130km to 560km with a mean of 380 km. For this specific analysis, the above described SNP filtering steps were carried out per species and not per cultivation line.

Second, we focused on the effect of cultivation on the genetic diversity within cultivated generations, estimated as expected heterozygosity *H*_e_. We hypothesized that genetic diversity would decrease with increasing numbers of generations in cultivation. To test this hypothesis, we calculated *H*_e_, using the unbiased estimator of Nei & Roychoudhury (1974), for each cultivation line and generation, taking allelic read counts as allele frequencies. To ensure comparability across species, we centered the *H*_e_ values per cultivation line and related it to the number of generations in cultivation, producer identity and their interaction as fixed explanatory variables and cultivation line as a random factor in a linear mixed effects model (Bates et al. 2015). When we visually inspected the data, *H*_e_ did not appear to change continuously across generation, but it showed an abrupt change between the wild plants and the first cultivated generation. We thus treated the generations in cultivation as a categorical variable in this model.

Next, we assessed how the different seed sourcing strategies of the two seed producers affected genetic diversity (*H*_e_). As a reminder, Producer 1 mixes seeds from multiple wild populations, while Producer 2 uses seeds from one population only, we expected the seeds from the Producer 1 to harbor higher genetic diversity. We tested this only for species where data from both seed producers were available (Table 1) and related the absolute values of *H*_e_ in each cultivation line and generation to species identity, generation and producer identity in the statistical models. The species and generation were included in the model to adjust for species identity and potential temporal trends across generations. For this analysis, the above described SNP filtering steps were carried out per species and not per cultivation line.

Finally, we tested whether the magnitudes of genetic differentiation and changes in genetic diversity caused by cultivation depended on species traits. We focused on two specific traits that we expected to have an effect, and that varied among our study species: seed dormancy and self-compatibility. The data on self-compatibility we obtained from the BIOLFLOR data base (Kühn et al. 2004) and additional literature (Table S1), and seed dormancy was estimated in a germination trial as the proportion of viable, but non-germinating seeds in a sample of the wild-collected (F0) seed (Table S1)). Other potentially relevant characteristics like harvest type (mechanic or hand) or frequency of harvest could not be used because we had this information only for seeds from Producer 1, where all except one species were mechanically harvested once per year. As response variables in these analyses, we used the genetic change between the wild and the first cultivated generation, because these two generations were available for most cultivation lines (Table 1). As a measure of genetic differentiation, we used the absolute *F*_ST_ value between F0 and F1. Changes in genetic diversity were quantified as Δ*H*_e_ = (*H*_eF1_ - *H*_eF0_)/*H*_eF0_, thus standardizing values across cultivation lines. A positive value indicated a gain and a negative value a loss of genetic diversity. We related these *F*_ST_ and Δ*H*_e_ values to the two plant traits in bivariate linear models with the *F*_ST_ or Δ*H*_e_ as response variable and self-compatibility and seed dormancy as explanatory variables.

## Results

The mean pairwise genetic differentiation between wild-collected seeds and seeds from the last generations across all cultivation lines of the 19 species was *F*_ST_ = 0.034 (range 0.002 - 0.071). When analyzed across all cultivation lines, *F*_ST_ significantly increased with the number of generations a population underwent cultivation (Table 2A, Figure 2A). The response of *F*_ST_ differed between the seeds from the two producers (Figure 2A): while there was an increase in the material from Producer 1, we found no changes in *F*_ST_ in the seed material from Producer 2. In the five species for which seeds from different regions were available, the absolute values of *F*_ST_ between wild-collected seeds and the last cultivated generation were much smaller (mean *F*_ST_ = 0.015, range 0.001 - 0.028) than the *F*_ST_ between wild populations from different regions (mean *F*_ST_ = 0.056, range 0.016 - 0.143, Figure 2B, Table 2B.)

**Table 2:** Results of different statistical models relating genetic changes in 20 cultivated wild plants to different aspects of the cultivation process or species traits. **(A)** The effects of generation in cultivation and seed producer identity on genetic diversity and genetic differentiation between wild-collected and cultivated plants (Fig 2A, 3A). Note that generation is a continuous variable in the model for genetic differentiation, but categorical in the model for genetic diversity (see the methods section for details). **(B)** Comparison between the genetic differentiation within cultivation lines and the natural genetic differentiation between regions (Fig 2B), for the five species where cultivation lines were available from two different regions (see Table 1). **(C)** Testing for differences in genetic diversity between the two seed producers (Fig 3B). In contrast to the model in Table 2A, this analysis was carried out only with species available from both seed producers. **(D)** The results of models testing the effects of self-compatibility and seed dormancy on changes in genetic diversity (Δ*H*_e_) and genetic differentiation (*F*_ST_) between wild-collected seeds (F0) and the first cultivated generation (F1) (Fig. 4 A,B). In multivariate models the terms are fitted sequentially. Significant values are in bold.

**(A)**
| Effect | Genetic differentiation ($F_{ST}$) | | | | Genetic diversity ($H_e$) | | | |
| --- | --- | --- | --- | --- | --- | --- | --- | --- |
|  | NumDF | DenDF | F | P | NumDF | DenDF | F | P |
| Generation | 1 | 50 | 4.53 | **0.038** | 4 | 71 | 7.40 | **<0.001** |
| Producer | 1 | 50 | 4.54 | **0.038** | 1 | 71 | 0.64 | 0.428 |
| Generation × Producer | 1 | 50 | 5.98 | **0.018** | 3 | 71 | 1.11 | 0.351 |

**(B)**
| Effect | Genetic differentiation ($F_{ST}$) | | | |
| --- | --- | --- | --- | --- |
|  | NumDF | DenDF | F | P |
| Within cultivation line vs.<br>between regions | 1 | 9 | 8.92 | **0.015** |

(C)
| Genetic diversity ($H_e$) | | | | |
| --- | --- | --- | --- | --- |
| Effect | NumDF | DenDF | F | P |
| Species | 1 | 19 | 5.94 | **0.003** |
| Generation | 1 | 19 | 0.23 | 0.639 |
| Producer | 1 | 19 | 26.51 | **<0.001** |

(D)
| Genetic differentiation ($F_{ST}$) | | | | | Change in genetic diversity ($\Delta H_e$) | | | |
| --- | --- | --- | --- | --- | --- | --- | --- | --- |
| between F0-F1 |  |  |  |  | between F0 and F1 |  |  |  |
| Effect | NumDF | DenDF | F | P | NumDF | DenDF | F | P |
| Self-compatibility | 1 | 16 | 16.55 | **<0.001** | 1 | 16 | 1.98 | **0.180** |
| Dormancy | 1 | 16 | 0.02 | 0.892 | 1 | 16 | 0.26 | 0.618 |

**Figure 2:**
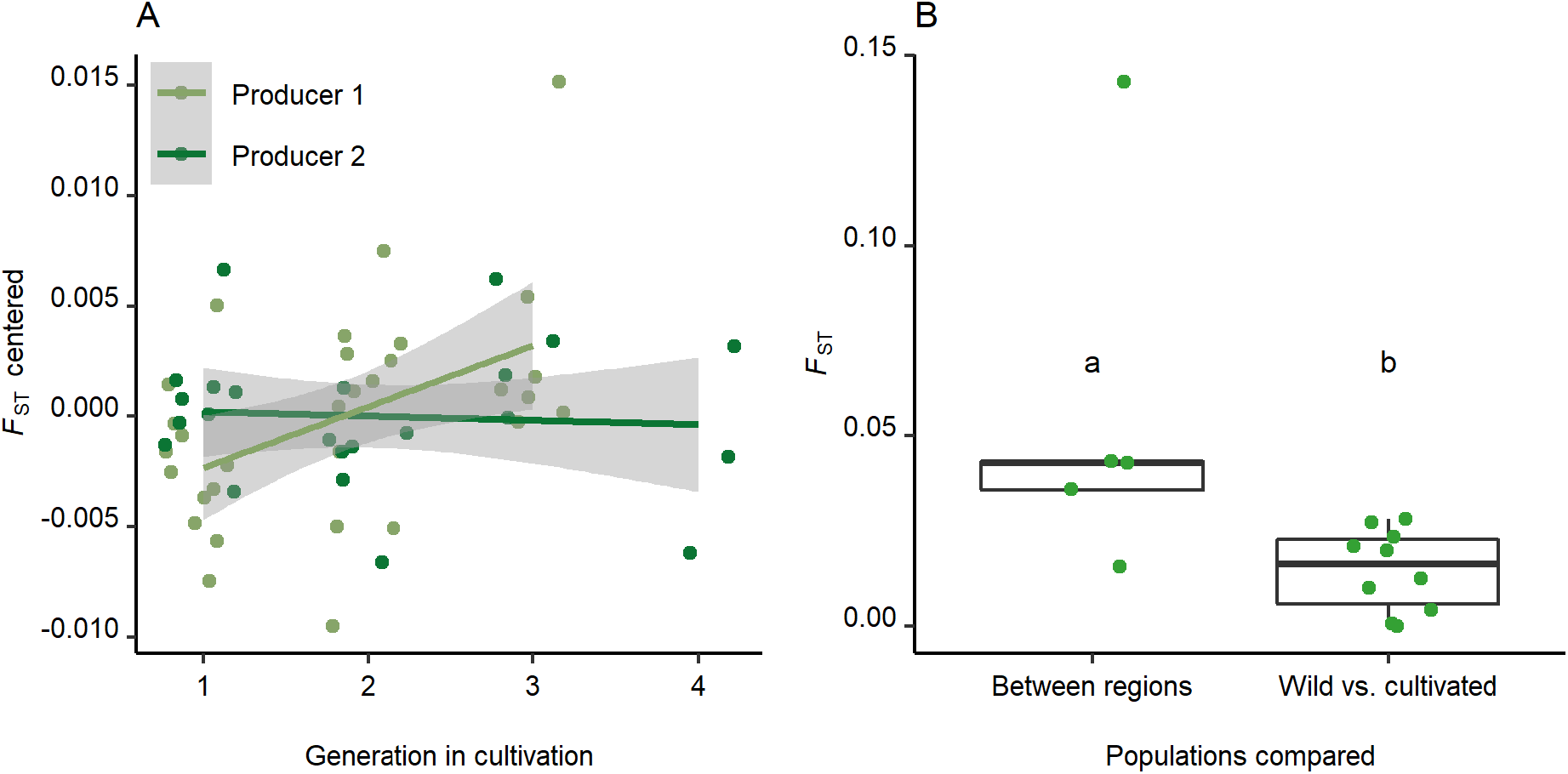
(A) the relationship between pairwise *F*_ST_ (centered to account for differences between species) between wild-collected seeds (F0) and later generations in cultivation across 19 wild plant species. Each point represents one generation of one independent cultivation line. The shaded area is the 95% confidence interval for predictions from a linear mixed effects model, Producer 1 (blue): *F*_(1,29)_=7.44, *P*=0.011, Producer 2 (red): F_(1,21)_=0.09; *P*=0.774. For the full model results, see Table 2A. (B) Comparison of pairwise *F*_ST_ –values between wild-collected seeds from the same species from two different regions (Between regions), and between wild-collected seeds and the respective last generation in cultivation (Wild vs. cultivated) for five species that were available from two regions; (see Table 2B for the model results). Each point represents a generation of an independent cultivation line. Letters above boxplots indicate significant differences.

**Figure 3:**
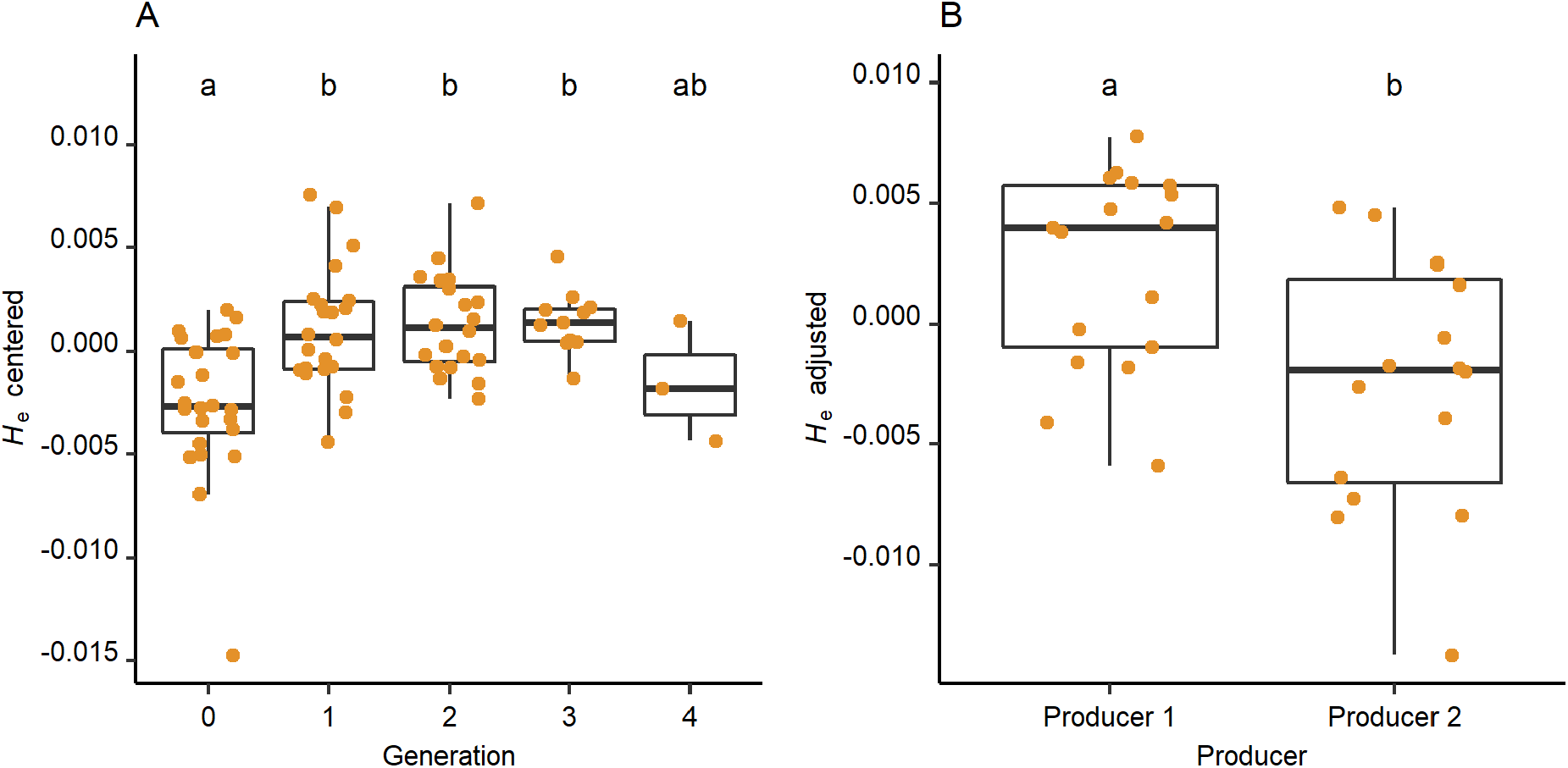
Effects of cultivation on genetic diversity. (A) Changes of (centered) gene diversity (*H*_e_) across generations in cultivation (see Table 2A for model results). (B) Comparison of genetic diversity between the two studied seed producers (see table 2C for model results). In (B) diversity was adjusted for species identity and generation in cultivation. Each point represents a generation of an independent cultivation line. Letters above boxplots indicate significant differences.

**Figure 4:**
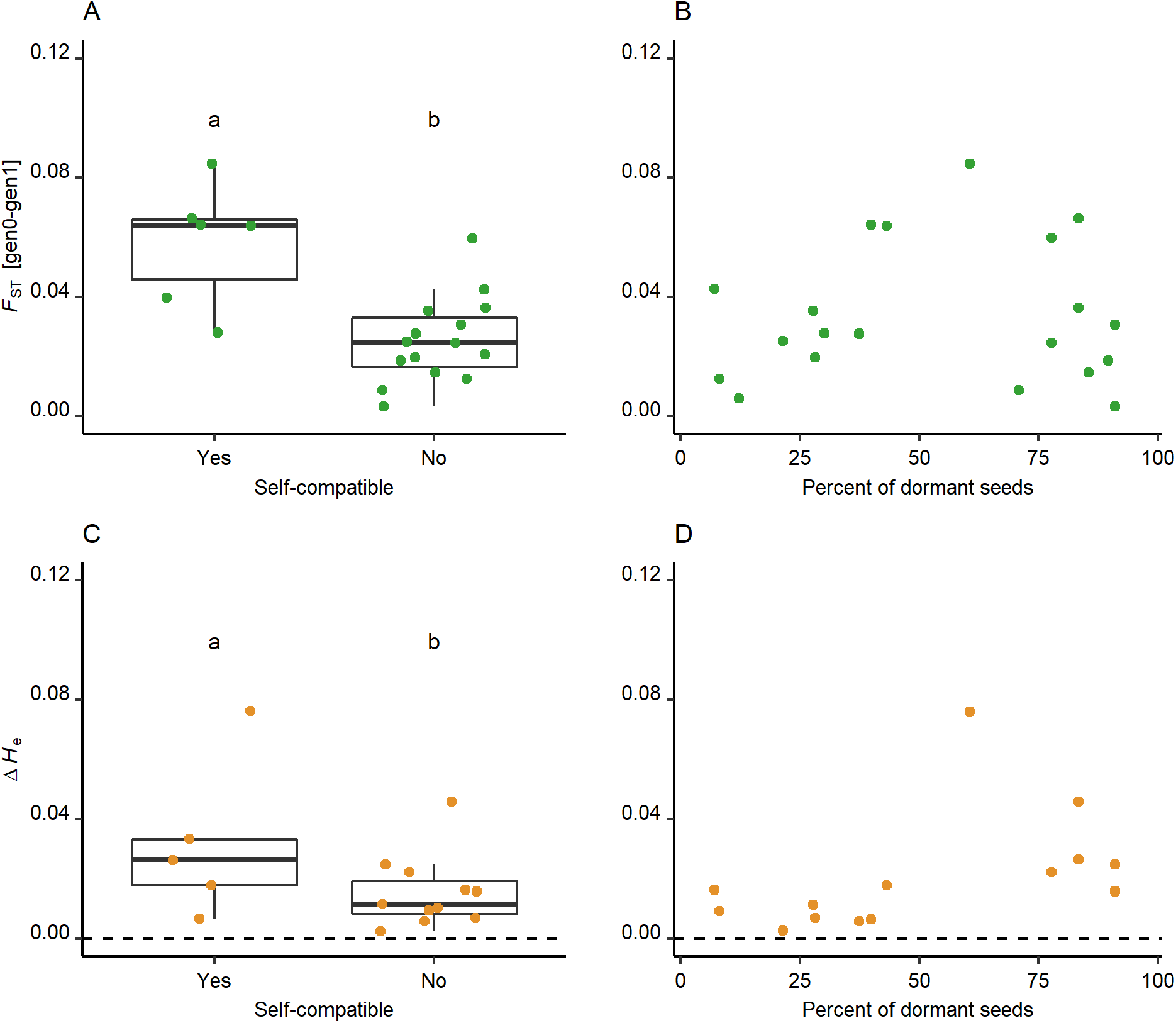
Relationships between the self-compatibility and seed dormancy of plants with the observed magnitudes of genetic changes during cultivation, specifically genetic differentiation (panels A and B) and changes in genetic diversity (panels C and D) between the wild-collected seeds (F0) and the first generation in cultivation (F1). Each point represents a species. The letters above boxplots indicate significant differences. For the full model results, see Table 2D.

The mean expected heterozygosity in our data set was *H*_e_ = 0.303 (range 0.282 - 0.337), and it was significantly lower in wild-collected seeds (F0) than in the subsequent generations F1, F2 and F3 (Table 2A, Figure 3A). Only the *H*_e_ in the last cultivated generation F4 did not significantly differ from wild seeds, probably because of small sample size of only three cultivation lines in the F4. While the effect of cultivation on *H*_e_ did not differ between the two seed producers (Table 2A), the seeds from Producer 1 harbored higher average *H*_e_ then the seeds from Producer 2 (Table 2C, Figure 3B).

The genetic differentiation (*F*_ST_) between wild-collected seeds (F0) and the first cultivated generation (F1) was significantly higher in self-compatible species compared to self-incompatible species (Figure 4A), with a similar, albeit not statistically significant, trend for changes in genetic diversity Δ*H*_e_ (Figure 4C). Seed dormancy was unrelated to genetic differentiation and genetic diversity (Figure 4B, D, Table 2D.)

## Discussion

Seed propagation for restoration is indispensable to cover the rising demand for native seeds, but the propagation process could change the genetic properties of the cultivated populations and impoverish their genetic diversity (Espeland et al. 2017). Here, we used molecular markers to test for such effects in 19 species cultivated by two different seed producers. We found that cultivated populations indeed genetically differed from the wild collected seeds, and this differentiation increased with the number of generations the populations spent in cultivation. Yet, the absolute size of the genetic differentiation due to cultivation was much smaller than genetic differentiation between natural populations from different geographic regions. The genetic diversity of cultivated seeds was even higher than that in wild collections, which suggests that cultivation, as done by certified producers in Germany and Austria, does not compromise the genetic quality of seed material.

### Genetic differentiation

The genetic differentiation between wild and cultivated plants increased with the time the populations spent in cultivation. Yet, this effect was rather small in comparison to the natural genetic differentiation between regions. Similarly, the few previous works that used molecular markers to study the cultivation effects on plant material propagated for ecosystem restoration also reported no or only minor genetic differentiation during cultivation in the majority of studied species (Nagel et al. 2019). In contrast, theory-based studies expected that cultivated plants could be substantially genetically differentiated from wild plants as a result of genetic drift and unintended selection (Espeland et al. 2017; Pedrini et al. 2020). This discrepancy between theoretical expectations and real data has multiple possible reasons. First, the expectations were formulated based on evolutionary theory supported by data from plant breeding and ex-situ collections in botanical gardens (Lauterbach et al. 2012). Yet, ex-situ collections have typically very small population sizes where genetic differences can quickly arise by genetic drift (Ensslin & Godefroid 2019; Rauschkolb et al. 2019). Propagation of plant material for ecological restoration typically involves large populations, where the effects of genetic drift are likely minimal. Second, we have been working with a system where the seed producers are aware of possible negative effects of cultivation on genetic properties of the population and they try to prevent them (Prasse et al. 2010; Krautzer et al. 2020; Bucharova et al. 2019). Such regulations may however not be present in all production systems (Jones 2013; Pizza et al. 2021). Third, the material we studied involved at maximum four, but mostly two or three cultivated generations. As the genetic differentiation was linearly increasing, cultivation for many more generations would likely lead to a more substantial genetic differentiation. Fourth, we worked with reduced-representation molecular markers, an excellent tool to identify population history and signs of genetic drift, but suboptimal for detecting adaptation (Lowry et al. 2016). The low genetic differentiation as detected in this study does not exclude selection at particular gene loci because adaptive traits may be coded by few genes that were not necessarily covered by our markers.

However, identifying loci under selection was outside the scope of our multi-species study as this would necessitate whole genome sequence data and reference genomes which are unavailable for our study species. A more realistic option to detect selective changes would thus be growing and phenotyping the plants in a common environment.

We did detect – although on a low absolute level – an increase of genetic differentiation across populations in cultivations in the material from Producer 1 but not from Producer 2. The producers differ in how they source the wild seed: Producer 1 starts with seeds from at least five natural populations, Producer 2 only with seed from one population (RegioZert 2019; Krautzer et al. 2020). The mixing of seeds from multiple populations within a source region, as performed by Producer 1, increases the genetic diversity (Boca et al. 2020). Higher gene diversity may provide a higher chance of subsequent evolution during cultivation. For example, the contribution of the individual source populations to the gene pool of the seed lot can shift during the propagation process (Basey St. Clair et al. 2020). The observed stronger differentiation between generations in the material of Producer 1 is thus likely caused by the more diverse starting material. However, diverse starting material also means a diverse seed lot, which is beneficial for restoration because it enhances the adaptive potential and reinforces the probability of the restoration success in a range of environments through the portfolio effect (Crowe & Parker 2008). The increase of genetic differentiation during cultivation in the seeds from Producer 1 thus likely does not mean that the production practice of this company is suboptimal, but rather that the benefits of enhanced genetic variability by population mixing is accompanied by an increased probability of minor evolutionary changes in response to the cultivation process.

### Genetic diversity

Genetic diversity was higher in the cultivated generations than in the wild-collected seeds, in the seed material from both seed producers. This contrasts with the common expectations that genetic diversity should decline because of genetic drift and repeated bottlenecks (Espeland et al. 2017; Kantar et al. 2017; Breed et al. 2018), as well as with data from ex-situ cultivations where genetic diversity often rapidly declines (Lauterbach et al. 2012; Ensslin et al. 2017). However, as pointed out above, ex-situ cultivations have small populations sizes, which makes them extremely vulnerable to genetic drift and subsequent loss of variability. Propagation of seeds for restoration typically involves large populations (Prasse et al. 2010), and the effect of drift is thus limited (Frankham et al. 2014).

The most likely reason for the observed increase of genetic diversity is enhanced outcrossing under cultivation. Under natural conditions, individuals that grow in close proximity have a higher chance of being closely related (Turner et al. 1982; Vekemans & Hardy 2004; Zeng et al. 2012), which enhances mating between relatives due to limited distances of pollen dispersal (Turner et al. 1982; Kunin 1993; Zeng et al. 2012). During all steps of seed propagation, however, seeds from individual plants are mixed, and neighbors are thus unlikely to be close relatives, even more so if multiple source populations are included. This, together with high population densities in the field, promotes outcrossing (Tong et al. 2020). As a result, cultivated seed lots likely contain more heterozygote genotypes and thus, harbor more genetic variability than wild collected seeds.

### Species traits and genetic changes

The magnitudes of genetic changes due to cultivation, estimated as the differences between wild seeds and the first cultivated generation, were significantly higher in self-compatible species than in obligate outcrossers for genetic differentiation and, as a trend, for genetic diversity. Wild populations of self-compatible species typically contain more homozygotes (Charlesworth 2006) than populations of outcrossing species. Consequently, the enhanced outcrossing during cultivation, means a stronger change for self-compatible species than for outcrossers. These results support our interpretation above that an increased outcrossing frequency in the large and homogenized cultivated populations is a major driver of the observed genetic changes during cultivation.

The percentage of dormant seeds in the source population was unrelated to the magnitudes of genetic changes. This surprised us. Loss of dormancy is among the best-documented evolutionary processes in the domestication of wild plants (Purugganan 2019), because genotypes with dormant seeds do not germinate readily after seeding and thus they do not produce any seeds that could be harvested. We expected that as dormant genotypes would get lost under cultivation in some species, these would experience more pronounced genetic changes. However, loss of dormant genotypes would be a result of selection. We possibly did not detect this effect due to methodic constraints, because reduced-representation sequencing is suboptimal for identify signatures of selection, which often affects only a small part of the genome and easily escapes detection (Lowry et al. 2016; Mckinney et al. 2017).

While our study provides valuable insight into potential propagation effects and suggests how species traits may affect the magnitudes of genetic change during seed production, the latter results should be interpreted with caution. For vast majority of the species, we had only one cultivation line, and we are thus not able to dissect between cultivation line – specific and species-specific patterns. Further, our study included only 19 species representing a narrow trait spectrum. To properly understand which species are more prone to genetic change during cultivation, we need to study more cultivation lines of more species representing a wider range of traits and production methods (van Kleunen et al. 2014).

### Implication for practice

Agricultural propagation of native plants for ecosystem restoration is mandatory to ensure sufficient amount of seed to achieve the ambitious targets set by The UN Decade for Ecosystem Restoration (Merritt & Dixon 2011). Yet, there have been concerns that propagation in production fields may compromise the genetic composition of wild seed and strongly reduce its genetic diversity (Espeland et al. 2017). Here we show that these concerns may be to a certain degree unwarranted, at least in the highly regulated seed production systems of Germany and Austria.

We indeed detected genetic differentiation between the wild and cultivated plants across species, but it was much smaller than the differentiation between wild populations of different regions. As the differentiation increased with the duration of cultivation within the 3-5 studied generations, it is possible that after additional generations the seeds from cultivation could become substantially different from the original seed sources (Pizza et al. 2021). We also cannot exclude selection on particular phenotypic traits due to cultivation and harvesting conditions. A cap on the maximum number of generations, e.g. five in the seed production we studied (Prasse et al. 2010), or in the Yellow Tag certification system in the USA (Young et al. 2003), therefore seems reasonable.

We found that genetic diversity during cultivation even increased, probably as a result of enhanced outcrossing rate in the production fields and, in some cases, mixing multiple source populations. This makes such farm-produced seeds especially suitable for restoration, because higher genetic diversity enhances adaptive potential and restoration success across a broad range of environments through the portfolio effect (Crowe & Parker 2008).

In summary, we show that propagation of native seeds for ecosystem restoration only moderately changes genetic composition of the cultivated seed lots, and that genetic diversity is not only maintained, but even increased. Yet, our approach did not allow to identify adaptive genetic variability and possible effects of unintended selection. Future research should attempt to close this gap through common-garden studies and transplant experiments that allow to test for changes in phenotypes and examine the adaptive significance of genetic changes.

## Acknowledgements

We are grateful to Saaten Zeller GmbH & Co. KG and to Wilhelm Graiss, HBLFA Raumberg-Gumpenstein, Austria, for providing seeds. This work was supported by DFG projects LA4038/2-1 to B. and DU404/14-1 to W.D.

## Author contribution

AB, OB and WD conceived the idea; MC with support from WD performed the laboratory analysis, MC analyzed the data with support from WD, CL and AB; MC wrote the first draft with significant input from AB and CL, all authors critically revised the manuscript.

## Data availability

Raw sequence data have been deposited in the European Nucleotide Archive (ENA, https://www.ebi.ac.uk/ena) under project accession number PRJEB47978 and individual sample accessions ERS7671398 - ERS7671620.

## Notes

### Competing Interest Statement

The authors have declared no competing interest.

